# Geometric classification of brain network dynamics via conic derivative discriminants

**DOI:** 10.1101/201905

**Authors:** Matthew F. Singh, Todd S. Braver, ShiNung Ching

**Author notes:** Corresponding Author. 600 S. Euclid Ave., 63110, St. Louis, MO, USA.

## Abstract

Over the past decade, pattern decoding techniques have granted neuroscientists improved anatomical specificity in mapping neural representations associated with function and cognition. Dynamical patterns are of particular interest, as evidenced by the proliferation and success of frequency domain methods that reveal structured spatiotemporal rhythmic brain activity. One drawback of such approaches, however, is the need to estimate spectral power, which limits the temporal resolution of classification. We propose an alternative method that enables classification of dynamical patterns with high temporal fidelity. The key feature of the method is a conversion of time-series into their temporal derivatives. By doing so, dynamically-coded information may be revealed in terms of geometric patterns in the phase space of the derivative signal. We derive a geometric classifier for this problem which simplifies into a straightforward calculation in terms of covariances. We demonstrate the relative advantages and disadvantages of the technique with simulated data and benchmark its performance with an EEG dataset of covert spatial attention. By mapping the weights anatomically we reveal a retinotopic organization of covert spatial attention. We especially highlight the ability of the method to provide strong group-level classification performance compared to existing benchmarks, while providing information that is synergistic to classical spectral-based techniques. The robustness and sensitivity of the method to noise is also examined relative to spectral-based techniques. The proposed classification technique enables decoding of dynamic patterns with high temporal resolution, performs favorably to benchmark methods, and facilitates anatomical inference.

## 1. Introduction

Understanding how different brain regions interact in task-dependent ways is a key goal of cognitive neuroscience. In this regard, a frequent aim of neural data analysis is to characterize the spatiotemporal patterns present within brain activity and, subsequently, to enable the association of such patterns with specific cognitive states. Perhaps the most classic of such approaches, certainly within the domain of electrophysiological brain recordings, is time-frequency analysis which involves the projection of data into the Fourier domain in order to examine synchronization, spectral power and cross-channel (i.e., network) relationships.

Recently, increased attention has been directed to more holistically describe neural recordings in terms of their underlying dynamics [1, 2, 3, 4]. Such approaches include methods for characterizing chaos and instability reflected in neural recordings via estimation of Lyapunov exponents [5, 6] and manifold reconstruction methods such as Takens embedding [7, 8], which has been used to infer directed information flow between brain regions without overt analysis of rhythmicity [9]. As dynamical systems-based characterizations, these approaches are unified in that interest is placed on directly characterizing the underlying vector field that ultimately governs the time-evolution of the observed recordings. The vector field describes the instantaneous evolution of dynamical systems i.e., the time-derivatives of its state variables. In this spirit, here we propose a technique for elucidating properties of the vector field that involves analysis not of the observed time-series, but rather their derivatives: the ‘velocity’ trajectories of neural activity. We use the term ‘velocity’ in the dynamical systems sense in which system states, such as a particular combination of cell voltages, are referred to as ‘positions’ within the state space and an observed sequence of ‘positions’ (states) forms a trajectory. The ‘velocity’ is thus considering how the system states change in time. It turns out that this rather straightforward transformation from considering system position to system velocity brings about several conceptual and practical advantages for classifying neural dynamics. Our approach is motivated in part by recent results in monotone dynamical systems theory that provide explicit characterizations of how the attractor landscape of a dynamical system relates to the geometry of its derivative phase flow ([10, 11, 12, 13]). In other words, by analyzing the observed velocity trajectories we can make certain deductions regarding the properties (e.g., the types and stability of attractors) of the underlying neural circuit dynamics.

Exploiting this notion, we postulate a paradigm for classifying different cognitive states by geometrically deducing and characterizing their corresponding velocity trajectories. As will be seen, this approach enjoys many advantageous statistical properties. In the particular case focused on in our paper, it reduces to an intuitive comparison of covariance geometry of the derivative time-series, which is amenable to classification using conic boundaries. We proceed to demonstrate the efficacy and ease of the proposed paradigm in a variety of examples, then apply it to an electroencephalogram (EEG) dataset [14] wherein we show its interpretability and performance in decoding covert spatial attention.

## 2. Results

### 2.1. Conic Analysis Reveals Underlying System Dynamics

We consider time-series (here, neural recordings) denoted as

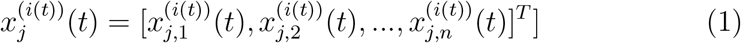

where *n* denotes the number of observed variables, 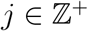 is a trial index and *i*(*t*) = 1, 2,…, *K* denotes a class label (e.g., a cognitive state). Our overall goal is to deduce the class label (at each time) by analyzing the derivatives of *x*_*j*_ (*t*).

The proposed method for neural data analysis and classification in this context consists of two steps. First, for each class (*i*), we obtain an estimate of the derivative time-series, 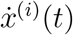. Second, we use generalized cones as a geometric classifier for individual points of the derivative time series. A generalized cone can be understood as an object in Euclidean space that is invariant to scaling: if a vector *v* is an element of a cone, then so is *αv* for any non-negative *α* ∈ ℝ^+^. Our approach will involve using such cones to partition Euclidean space into sets of derivatives attainable only by specific classes. Thus, the cone is a fundamentally directional object and that can reveal how a system transitions between states (but not necessarily the speed at which these transitions occur).

Our subsequent results will demonstrate that this approach complements spectral analysis (e.g., assessment of oscillatory power), which, while an assay of dynamics, does not explicitly include directional information. More specifically, unlike many existing methods for spatiotemporal classification, we do not operate upon summary parameters such as power within a specific frequency band ([15], [16]). Instead, it is the individual data points of the derivative time series that are used as the basis for classification since these contain inherent spatiotemporal information. In this way, our classification approach can capture the instantaneous rules, (i.e., the dynamics), governing how the (class-specific) time-series evolve. This approach is particularly favorable as it reduces a spatio-temporal classification problem in the original time series to a spatial-only problem in the derivative time-series. As we demonstrate later, this reduction significantly aids analysis and interpretation.

The motivation for such an approach stems from recent results in the theory of smooth monotone dynamical systems. For our purposes, a smooth dynamical system consists of a state space (here, we consider this to be ℝ^*n*^) coupled with a phase flow, or vector field that describes, for each state, how the system evolves forwards (and, backwards) in time. Recent research has linked geometric knowledge about the vector field (i.e., the derivatives along system trajectories or paths) to the asymptotic behavior of the system state variables [10],[11],[12],[13]. Such characterizations have been made possible by formally studying the extent to which the derivatives along a path may be confined within generalized cones. Although a detailed discussion of monotone dynamics is beyond our current scope (see e.g., [17]), these recent advances motivate our use of cones for classification in derivative space.

Thus, in as much as spatial patterns are useful for classification, the derivatives can provide us crucial information regarding the underlying system dynamics. Perhaps an intuitive analogy is that of vertical ascent for two sorts of aircraft: (fixed-wing) airplanes and helicopters. While helicopters can make a direct vertical translation, airplanes require strong lateral velocity before any vertical motion is obtainable. In contrast, reaching a point just above by airplane requires a complex velocity trajectory. Observing this velocity trajectory is thus highly informative in deducing the vehicle in question. Aerodynamic constraints also greatly constrain the turning radius of an airplane. As a result, the set of obtainable velocities for an airplane relative to its body-frame differs greatly from that of a helicopter, despite both being able to reach any point in airspace. Similarly, neural circuits are seemingly able to produce a wide variety of spatial activity patterns, although the set of attainable derivatives may be more limited. For example, in Figure 1 we consider two different toy networks, each composed of three continuous, recurrent Hopfield-model neurons ([18], see Methods; Fig 1A, B). The two networks (systems) feature a common connection scheme, but differ in their internal dynamics. In the second system (“System B”) every cell exhibits greater decay (in neurosciece parlance, leak conductance) (1B). The systems produce qualitatively similar trajectories wherein all initial conditions lead to a damped oscillatory response for both cases (1C,E). Despite this similarity, the modeled systems differ in the oscillatory envelope of their response (1F). In the ideal case of noise-free uninterrupted observations, the trajectories of these systems are easily discriminable in the original phase space (1E). However, trial-based designs that are the primary workhorse of experimental neuroscience recordings (e.g., single- or multiunit, EEG, fMRI) generate large numbers of short duration observations, in which the variability of initial states present at the trial start can result in significant entanglement of activation trajectories. When we replicate these conditions for the two simulated networks, their recordings become inextricable in the phase-space of cell activity (1C). However, the systems remain easily separable in the phase space of their derivative signals (1D). As the derivative space is indicative of instantaneous changes in activity, dynamical features of the system are captured in individual observations, so the systems may be differentiated regardless of observation length or initial conditions. Of course, this classification can be performed to a certain degree via direct methods (e.g., linearization) if the full analytical model is already known. Our goal, in contrast, is to make the same dynamical inferences (i.e., about the structure and equations governing the system) based only upon observation of the activity. Using our proposed approach, we observe that derivatives are invariant within a particular directional region (1D). Based upon this invariance, we can directly surmise that System A (1A) cannot directly move across the Cell 1 axis as its derivatives can never point in that direction (1). Instead, its derivative space suggests System A could only traverse the space by moving indirectly in that direction, just as physical constraints prevent vertical translation of a fixed-wing aircraft. One possible solution, is for System A to generate a series of oscillations leading to traversal of the space along the Cell 2 axis, which is of course consistent with the actual dynamics (1E,F). In contrast, derivatives of System B are rarely orthogonal to the Cell 2 axis (1D) and the corresponding prediction holds that System B traverses the Cell 2 axis more directly (with fewer oscillations) than System A (1E,F).

To operationalize the above analysis, we propose a classifier to discriminate systems based upon their directional derivatives. Most classifiers used to analyze neural data, such as linear support vector machines ([19]), assume that the classes differ in some combination of their univariate means and thus form linear boundaries ([20]). For the case of derivatives, this assumption is not valid. In fact, a well-known result in dynamical systems theory, which we rely upon, is that the derivatives of a smooth, bounded dynamical system must be balanced (e.g. [11]): over long scales the mean derivative is always zero. Otherwise, the system would grow infinitely along the direction of the mean derivative. In the case of a neural system, the mean change in the measured activity variable (e.g., voltage) must be zero over a sufficiently long recording, lest the activity grow infinitely. As the mean derivative is thus irrelevant, the motivating results in dynamical systems theory concern invariance along cones. Due to scale-invariance, conic boundaries discriminate between classes based upon the directional components of the data, rather than magnitude features (such as speed of evolution), and are always centered at zero. With very mild assumptions upon the way derivatives are distributed for each class, we can derive a Bayes-optimal classifier for directional data (see Materials and Methods)which reduces to a comparison of covariance matrices. The boundaries of this classifier form a quadratic cone.

For an unlabeled time-series *x*^(*i*(*t*))^(*t*) we define our quadratic-cone classifier as:

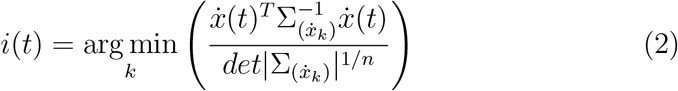

Here, 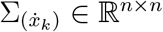 is the covariance matrix assoicated with the derivative time-series of class *k*. At an implementation level, this covariance can be estimate numerically by collecting labeled training data from each class. Thus, the classifier operates on a point-wise basis: assigning a class label to each individual data-point of the derivative time-series. In the two class case with class labels ±1 we describe the cone in terms of a single matrix *P*:

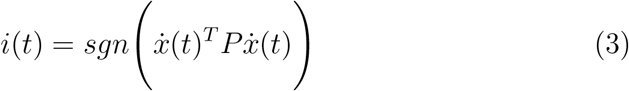

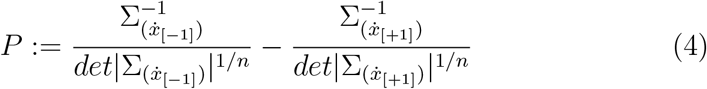

We denote the induced cone as *C*(*P*):= {*y* ∈ ℝ^*n*^|*y*^*T*^*Py* ≤ 0}. The decoding rule may be restated in terms of the cone *C*(*P*) as 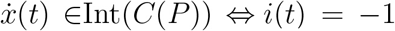 with Int indicating the interior (i.e. when the inequality is strict). We present further information concerning practical and efficient calculations of these terms and the case of singular covariances (i.e. fewer samples than channels) in Materials and Methods. Formally, we use the term ‘derivative’ in the sense of backward approximation by Newton’s difference quotient, however the scale-invariance property of cones makes such formalism unnecessary and the backward derivative thus becomes the difference time series: 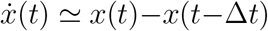. In general, we have had success implementing the derivative in this form without further preprocessing, even though an extensive literature exists concerning more sophisticated methods of numerical differentiation (i.e. [21]).

### 2.2. Conic Geometry Blindly Identifies Upstream Network States

To test the degree to which conic-invariance describes changes in functional structure ([22]), we simulated the case in which an experimenter tries to study a complex neural network but may only access a small number of outputs (Fig. 2A). We formalized this scenario by simulating a chaotic ([23]) system’s effects on a downstream recurrent network of four bursting Hindmarsch-Rose model neurons ([24]) with recordings only available for the first three cells (Fig. 2B,C). Cells are either excitatory (green) or inhibitory (red). The fourth, unobserved, inhibitory neuron forms the main recipient of upstream input and is normally quiescent save when activated by the upstream component. When the upstream component is in the “off” state, the network acts as a delayed negative feedback loop. When the upstream component is in the “on” state, a new indirect inhibitory path is opened via Cell 4. Thus, by altering the activity of a mediating cell, the upstream component’s activation dictates possible interactions in the downstream network.

**Fig 1.**
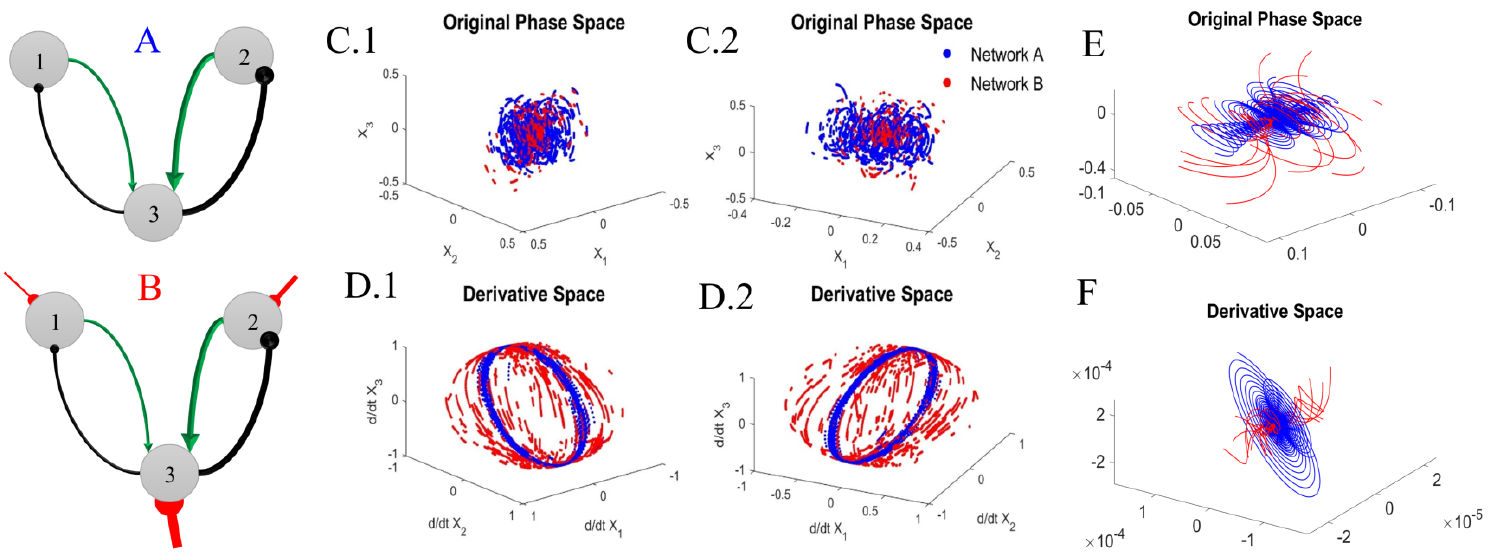
The two variants of a recurrent continuous neural network (A,B). Line widths indicate connection strength and green/black indicates excitatory/inhibitory. The red markers in network B indicate a greater decay rate. Observed trajectories do not differentiate between generative systems in the original state space (C.1, C.2), but are readily distinguishable in the derivative space and its projection onto a spherical surface (D.1, D.2). Random intervals of each orbit are plotted for 150 initial conditions and two views (view 1: C.1,D.1; view 2: C.2,D.2) with derivatives projected onto ∂*S*^*n*−1^. In the original phase space (E) full trajectories of both systems approach a focus, however, they differ in the approach envelope which separates derivatives (F).

In order to blindly determine the upstream component’s activity state we performed unsupervised clustering of downstream neural voltage. Using the three observed neurons we performed 2-means clustering ([25]) for either the spectral power of the original voltage trace (*V*(*t*)) or, for conic analysis, the covariance of the derivative voltage traces 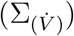. Spectral power and covariances were calculated over equal-length non-overlapping time windows. Using the k-means algorithm ([25]) produces both a set of centroids (cluster means) and labels assigning each data point to the cluster to which it is closest. For the spectral classification we directly used the labels assigned to each time bin by k-means as the training class. For conic classification we discarded the original labels for each time bin and instead generated new labels through our proposed conic method. The two covariance centroids were treated as class covariances and we then classified each time point according to (2). Because the conic classifier provides instantaneous class labels, we assigned whole-bin classes post hoc in a winner-take-all paradigm based on the

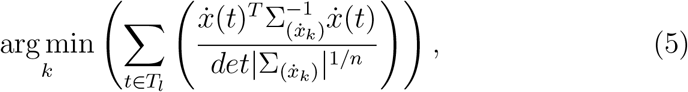

where *T*_*l*_ denotes the *l*^*th*^ bin.

We repeated this simulation/analysis sequence while varying simulation parameters for a total of 136 simulations per case (see Materials and Methods). The parameters of interest were the intrinsic system noise (a Brownian motion), measurement noise, coupling strength with the upstream component, and the window length for each bin. Results demonstrate a general advantage of conic classification over spectral clustering (Figure 2D). This relationship was most pronounced for strong coupling strength or low measurement noise. Unlike conic decoding, spectral decoding was largely unaffected by the white-noise added to the recorded voltages. Spectral performance exceeded conic decoding in extreme cases of measurement noise, which is well known to degrade estimation of derivative time series. However, both methods were resistant to intrinsic noise within the system. These two noise varieties (intrinsic vs. measurement) had equal variance ranges, but unlike measurement noise, intrinsic noise interacts with system states. For instance, Brownian motion in the voltage variable has less impact during the fall of an action potential than near the firing threshold, so this differential sensitivity may preserve those portions of dynamics to which the conic method is most sensitive. We conclude that for most parameter choices within this simulation, the proposed conic decoding method exceeded spectral classification of upstream system states but is susceptible to idealized measurement noise. However, we later demonstrate that the conic method is actually resistant to many (realistic) sources of experimental noise and artifact (see Discussion).

### 2.3. The Conic Method Produces Instantaneous Decoding of Covert Spatial Attention

In the next experiment, we consider the method’s application to an EEG dataset ([26],[14]) recorded during a spatial covert-orienting task. During the task a briefly presented central cue indicated where a subsequent task-relevant target would appear with 80% accuracy. The central cues had near identical visuo-spatial features and subjects were instructed to focus upon the cue (see Materials and Methods). This design feature implies that the putative neural dynamics associated with particular orientations correspond to covert spatial attention rather than simply the direction of gaze. Previous results using this dataset have demonstrated that the spatial distribution of alpha power (across channels) may be used to determine to which location subjects are covertly orienting. These previous analyses ([26]) were performed pair-wise using L1-regularized logistic regression for alpha-band power in parieto-occipital electrodes and the authors presented the best pair accuracy for each subject (74.6 ± 2.3%) as well as the number of significantly classified pairs per subject: (3.5 ± 2.67) out of a total of 15. In our comparison, we use all trials with delays of at least 1300ms and only consider data after the first 250ms post-cue. We consider the same metrics to those previously reported for comparison ([14]) and all reported accuracies for the current analyses are based upon leave-one-out cross-validation.

**Fig 2.**
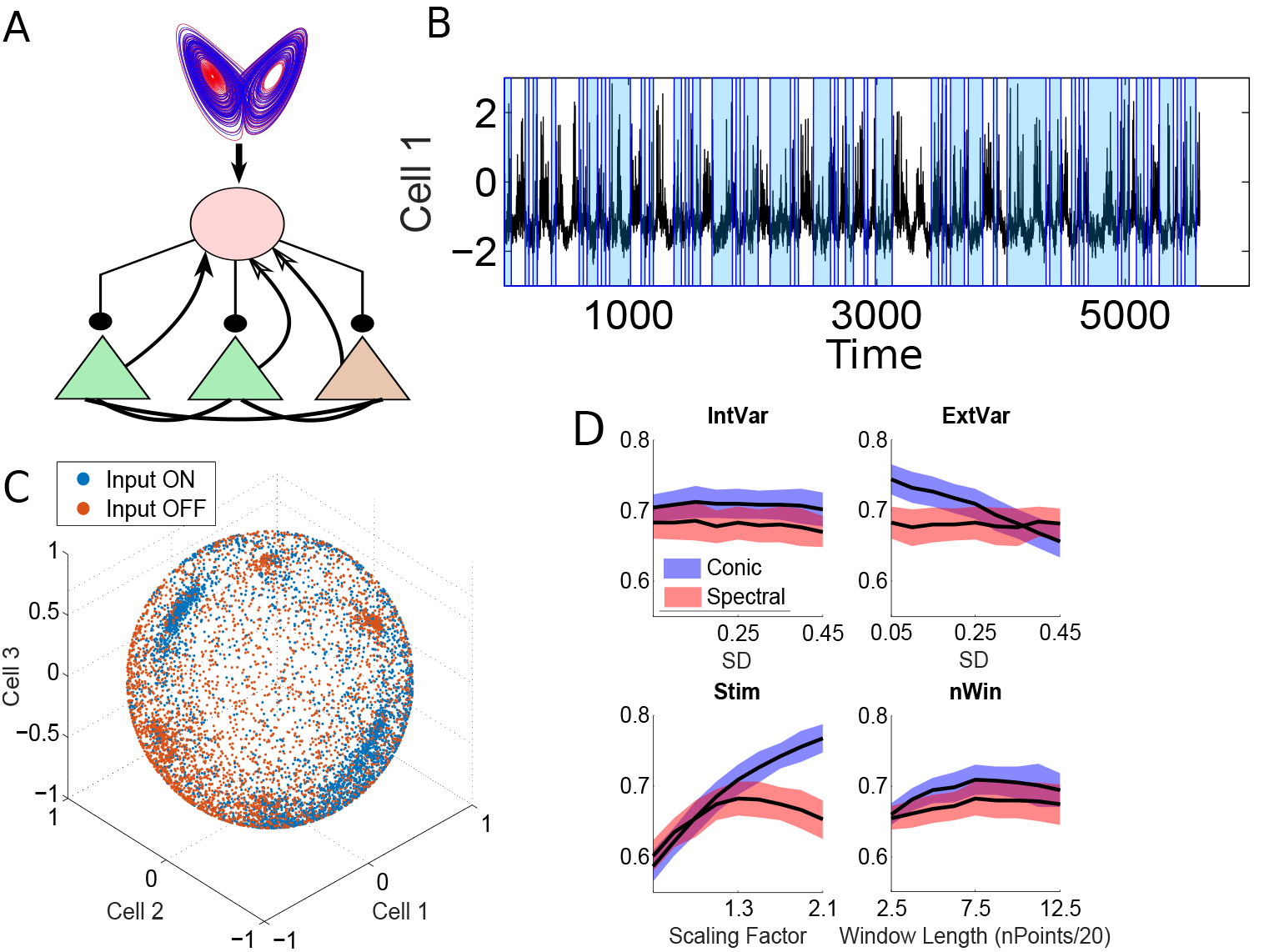
Simulated comparison of blind conic vs. spectral classification. A) The system consists of an upstream chaotic attractor which provides binary downstream input to a two-layer recurrent system. Green cells are excitatory and red inhibitory. B) Example simulated voltages for the the three cells in the bottom layer using median model parameters. Blue bars indicate when the upstream input was “On”. C) The state of the upstream system is easily determinable by plotting the voltages in the normalized derivative space. Voltage traces are in black and blue shading indicates periods in which the upstream system was “on”. D) Percent correct by decoding method and simulation parameters. Neither method was affected by the level of intrinsic system noise (IntVar), but conic decoding suffered with the addition of measurement noise (ExtVar). Conic decoding accuracy continued to increase with the upstream coupling strength (Stim), while spectral decoding showed a concave relationship. Both methods performed slightly better for moderate bin sizes (nWin) over very small ones but further increases did not improve performance. The Shaded regions give standard deviations (n=136).

#### 2.3.1. Group-Level Conic Performance Exceeds the Limits of alpha-band Classification

In the case of pairwise classification, current results appear favorably [Figure 3 A,B] to those previously published ([14]). In their analysis, Schmidt and colleagues ([14]) provide two metrics: the number of significantly discriminated pairs and the maximal pairwise accuracy. We emphasize comparisons in terms of number of significant pairs as these take into account more subtle spatial comparisons whereas the previously reported maximal pairwise accuracy corresponded to spatially opposite locations for seven out of eight participants in [14]. Results demonstrate that the conic classifier is, on average, superior to *α*-band based decoding in terms of number of significant pairwise classifications (*t*(15) = 3.558, *p* = .004, 2-*tailed*), conic pairs: 10.25±1.8. However, the conic advantage in maximal pairwise accuracy was insignificant (*t*(15) = .572, *p* = .577, 2-*tailed*), conic maximal accuracy 76.4 ± 3.2%.

#### 2.3.2. Conic and Spectral Decoding Form Complementary Techniques

As with traditional approaches applied to the dataset ([14]), conic decoding exhibits variable performance across subjects. Interestingly, however, conic and alpha-band decoding performance across individuals tended to be negatively correlated in terms of the number of significant pairs per subject (Figure 3C). Due to the small sample size and suspected outlier (Subject 8), we performed a rank-order test with the outlier and parametric test both with and without the outlier. Using the full data with rank-order correlation demonstrated a negative relationship between conic and spectral performance (*Kendall’s τ*(6) = −.5879, *p* = .0738, 2-*tailed*). The parametric relationship with the full data did not reach significance due to the low sample size (*r*(6) = −.355, *p* = .3882, 2-*tailed*). This relationship greatly increases (*r* = − .940, *p*(5) = .0016, 2-*tailed*) when removing subject 8 for whom no pairs were significantly classified by the conic method. However, the low sample size makes it difficult to determine whether the suspected outlier does in fact represent anomalous data or just the extreme poor end of conic classification ability. Thus, while the small sample size interferes with statistical inferences, results suggest that not only are these decoding methods complementary, but also that many subjects exhibiting poor spectral-based decoding may actually possess strong conic classification. In summary, the proposed conic method and spectral decoding may form complementary approaches and subjects insensitive to one technique may benefit from the other.

**Fig 3.**
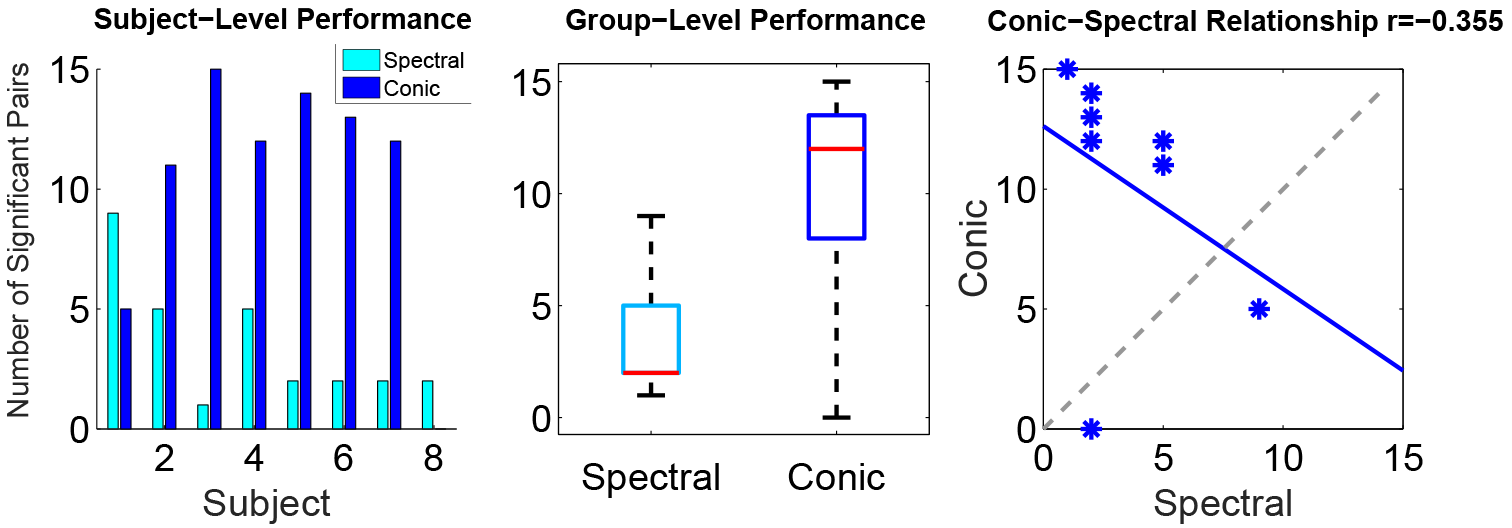
Comparison of conic and alpha-band based decoding methodsin terms of number of significantly decoded location-pairs. A) Both decoding methods possess large inter-subject variation in decoding performance. B) At the group-level conic decoding outperforms spectral. C) At the individual-level the techniques are negatively correlated and thus form complementary methods.

#### 2.3.3. Instantaneous Decoding Tracks the Evolution of Cognitive States

The conic-geometry of these instantaneous changes is easily visible for the 2-class case (i.e. lower left vs. upper locations) when projecting the derivative time series into the eigen-coordinates (see Materials and Methods) with the largest eigenvalues (Figure4A,B). These may be thought of as ‘principal components’, but as they relate the differential activation between classes they also possess sign information. Further, the absolute magnitude of each eigenvalue indicates how strongly selective it is for the class associated with its sign (Figure 4B). The time-series of these components indicate that the information differentiating between classes is associated with increased dynamics of certain spatial patterns (the eigenvectors). Critically however, these dynamics are not necessarily associated with consistent increases in particular frequency bands (Figure 4B). Point-wise classifier performance may be visualized by projecting data into the plane of positive and negative eigenlengths (see Materials and Methods) which indicates the combined ‘magnitude’ of dynamics associated with each class (Figure 4C).

#### 2.3.4. Conic Weights Generate Retinotopic Maps of Covert Attention

The conic method also appears to replicate previously published results showing that spatial aspects covert-orienting of covert orienting are most prominent of posterior electrodes. In fact, spatial location is highly prominent in the conic weights (see Materials and Methods) and, in this case, provides anatomical localization of dynamical information (Figure 5) without requiring search-light type analyses ([27]). To determine whether conic weights provide anatomically meaningful information we performed a spatial mapping of the weights for each cone-generating matrix. We defined the map for each condition in terms of the contrast ‘location x’ vs. the average hemifield (3 locations) opposite ‘x’. We averaged the covariances across subjects for viewing purposes (group level), but in most cases similar results held at the individual subject level as well. As a matrix, the conic classifier is most naturally visualized as a weighted graph. However, to produce a spatial map we assigned each channel a weight based upon its contribution to each of the matrix’s eigenvectors (see Materials and Methods). We find that the regions representing the covertly attended location experience decreases in the amplitude of derivatives (negative weights) while regions representing the opposite spatial location experience relative increases in the amplitude of derivatives (positive weights) (Figure 5). Thus, covert attention may be associated with neural activity whose variability (as distinct from variance) is more restricted in regions representing the covertly attended location.

**Fig 4.**
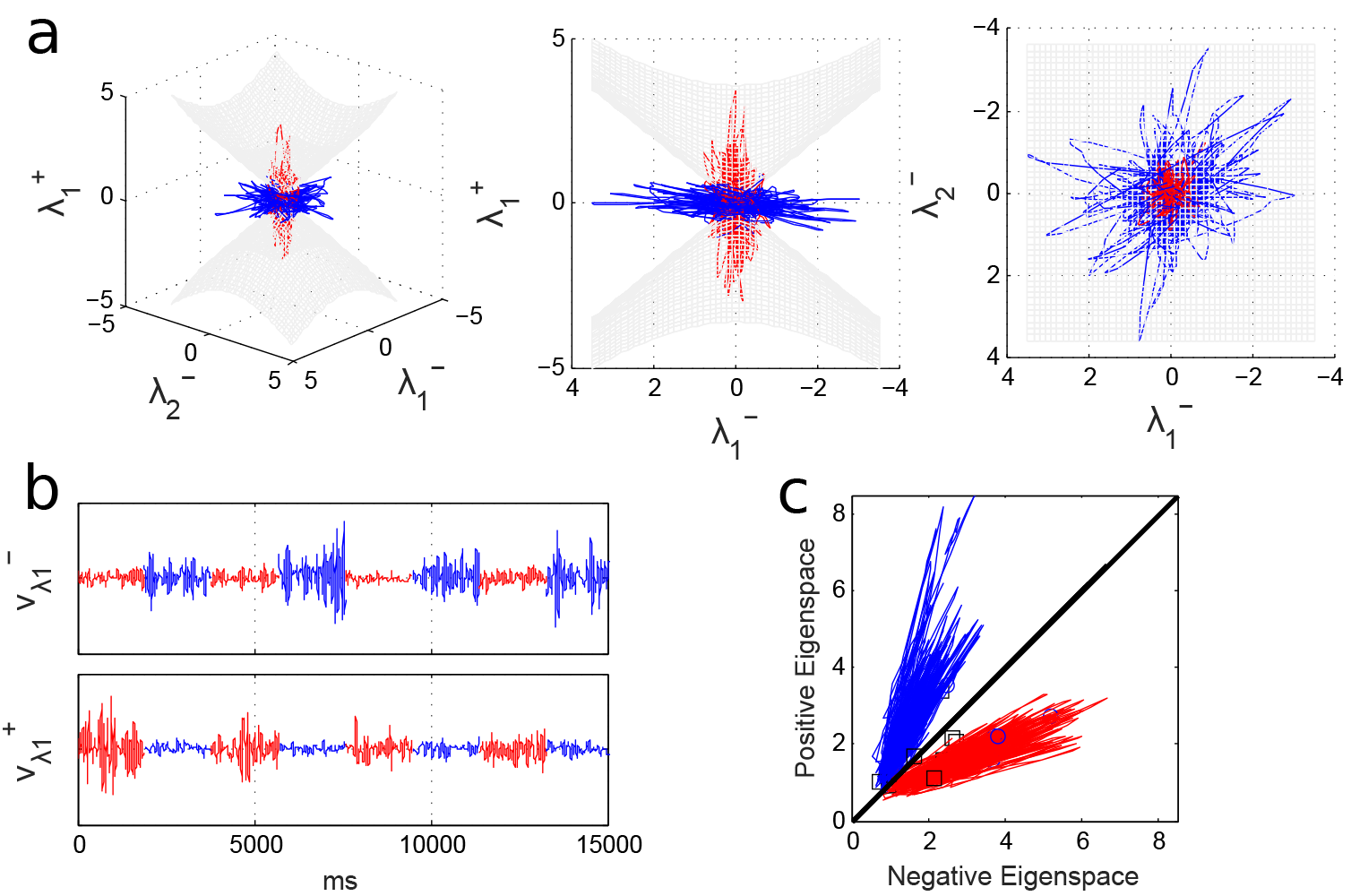
Conic Geometry. A) Plotting the first 2 negative components and the first positive component in derivative space reveals conic geometry. Data is displayed from a variety of viewpoints. B) Plotting the primary positive and negative eigencomponents of the cone for a 2-location contrast demonstrates that these components increase derivative amplitude during the corresponding cognitive state. C) Two dimensional projections onto the length in the “positive” and “negative” eigenspaces for the discriminative cone reveals robust classification of individual observations lasting only a few milliseconds.

**Fig 5.**
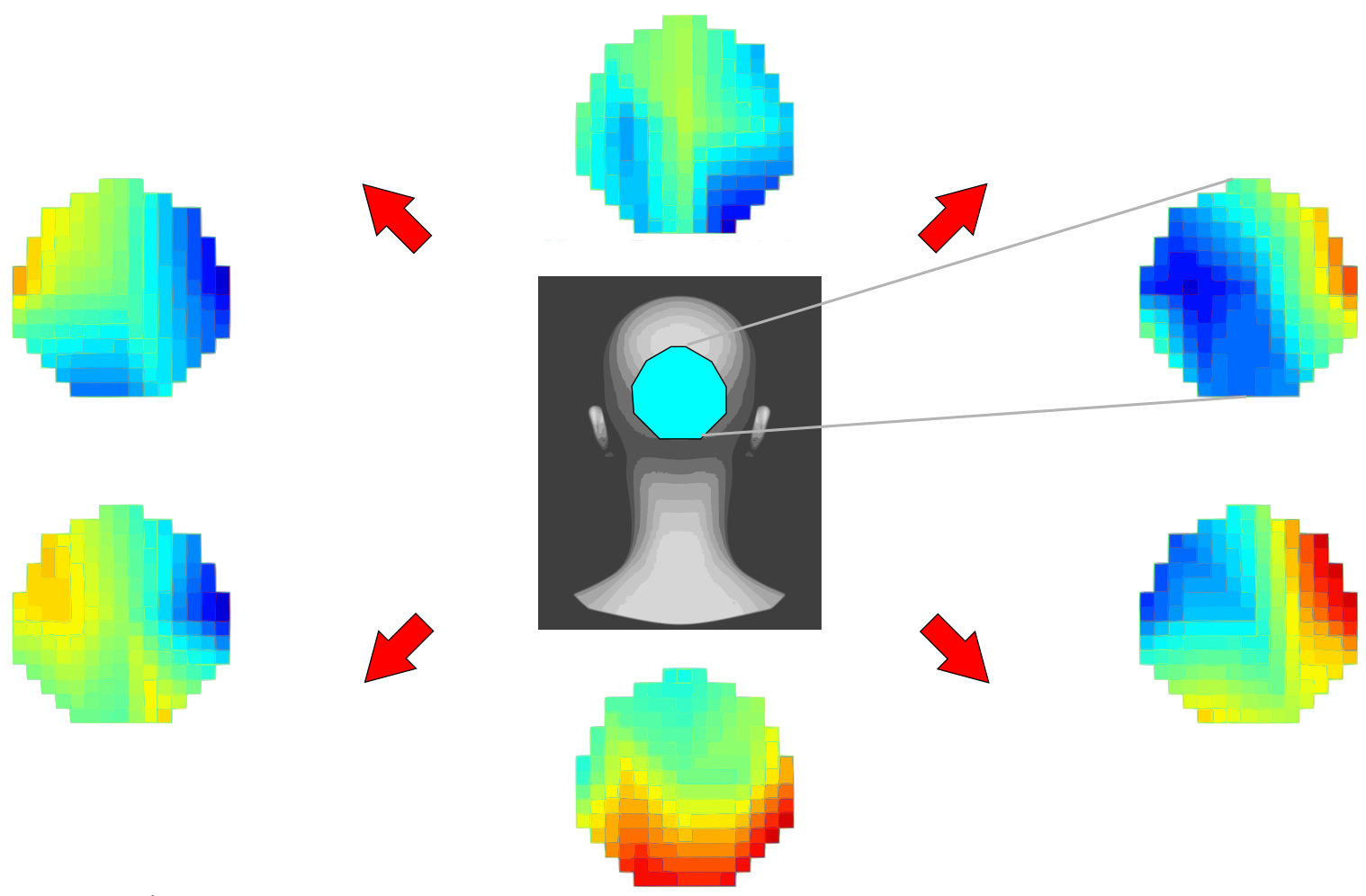
Anatomical weights distributions. Inverting conic weights to anatomically map classifier features produces a retinotopic map of covert spatial attention in posterior channels. Center panel indicates the corresponding region of head space (posterior coronal) while radial panels denote the weight mapping for subjects attending to that location vs. opposite locations with hot colors indicating “increased” dynamical instability.

## 3. Discussion

We have introduced a novel method for performing classification of spatiotemporal signals. The key innovative feature of the approach is the use of the derivative rather than the original time-series as the basis for classification. By using the derivative time-series, our conic approach to classification is sensitive to temporal information even though the classification method does not explicitly consider time as a variable. After computing the derivative time series, the method proceeds identically to static classification methods, with each time point of the derivative time series corresponding to one sample. From a dynamical-perspective the conic approach emphasizes decoding based upon the vector field of a system rather than the time course of trajectories. For a bounded signal, the distribution of derivatives must center about the origin for any suitably long segment, rendering linear decoding of the derivative space infeasible. However, this balancing property makes derivative space particularly well-suited for conic boundaries, as information is concentrated in the direction of points rather than their magnitude. Consequently these can be well-described according to properties of limit-sets for cone-invariant systems ([10],[11],[12], [13]). We have given one example of an explicit form for conic classification which bears some similarity to a special case of quadratic discriminant analysis. However, unlike conventional quadratic discriminants, the balancing properties of derivatives and scale-invariant property of cones enable a natural explicit form for the n-class boundaries. These properties also lead to a number of invariance properties that cancel certain forms of noise such as scalar-multiplicative noise (i.e. amplifier noise), multivariate box-wave noise and low frequency noise (as elaborated on below). The form we provide relies exclusively upon the covariance matrices for each class’s derivative time series and is essentially a comparison of the degree to which their derivative covariance matrices differ.

### 3.1. Invariant Properties Promote Robust Classification

The proposed approach yields a number of fruitful properties that minimize the contaminating effects of variance due to both ‘noise’/artifact and within-subject variance. The conic classifier is robust to three sorts of noise: (slow) scalar multiplication, temporal warping, and multivariate box noise. The first two relate to the scale-invariant properties of cones. The first case results from simply differentiating the product of the original time series *x*_*k*_(*t*) and a scalar function *f* (*t*). When *f* is slow, we have 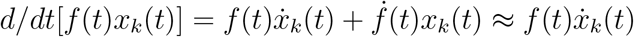. Due to the scale invariance of cones, 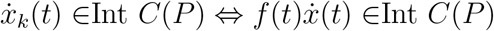 when *f*(*t*) ≠ 0. This case is especially relevant for multiplicative noise in which a multivariate signal is multiplied by an unknown scalar function, as could be the case for amplifier noise or changes in reference electrode conductance for EEG recordings. However, while the scaling will not affect how datapoints are labeled for a fixed classifier, it is possible that bias in the scaling function’s magnitude can affect the calculated covariance, hence the corresponding classifier. Normalizing individual data points of 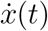 can remove this effect, although this procedure also can change the calculated covariance.

When sampling rates are high so that the time-series are relatively smooth, the conic method is also invariant to temporal scaling. More formally, we define a class invariant interval as a continuous interval *τ*: = [*T*_1_, *T*_2_] ⊂ ℝ+ with*t*_1_, *t*_2_ ∈ *τ* ⟹ *i*(*t*_1_) = *i*(*t*_2_). Let *y*(*t* ∈ *τ*) = *x*(*f* (*t*)) for any strictly positive monotone *f*: *τ* ⟶ *τ* in which *τ* is a nonnegative class-invariant section of the time series. Then 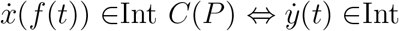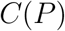. This case follows immediately by applying the chain rule and noticing that strict monotonicity implies 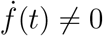. As the conic method is invariant to scalar multiplication, so too is it invariant to temporal “stretching” which corresponds to a conserved order of events despite irregularity in the precise timing. This problem of temporal “stretching” is referred to as dynamic time warping (DTW), and has been discussed by many previous authors, including in the use of DTW on derivative time series for classification ([28],[29],[30]). In general, dynamic time warping algorithms give a similarity measure between time series evolving at different rates by comparing the order of events between time-series rather than on a point-by-point basis. The recently proposed derivative-DTW variants operate similarly, but on the time series of first and second derivatives. Thus, both DTW and the currently proposed method are able to perform classification on warped time series. Unlike DTW, however, our approach does not explicitly consider the order of events, but rather operates on the set of realizable changes (the derivative geometry) and is thus applicable to systems in which different initial conditions also produce different orders of events. For example, in the Lorenz attractor ([23]), different initial conditions not only affect how long a trajectory stays within one of the attractor’s two “loops”, but also how many cycles it completes before exiting. The resultant time series from very similar initial conditions within the same system thus produce time series whose order of events differ greatly, despite generating identical state-space geometries after sufficient time. As with the previous case of multiplication by a scalar function, temporal warping will not affect how data points are classified for a fixed cone, but it can affect the calculated covariance, hence the classifier. As before, this effect can be removed by normalizing individual data points of the derivative time series.

The method is also resistant to additive multivariate box noise. This follows simply from the fact that the box function has derivative zero everywhere save at a finite number of points (corresponding to steps) and thus has minimal influence on the derivative time series. When the sampling rate is sufficiently high, many slow signals such as those induced by motion artifact may be approximated via a series of box functions, making conic decoding robust to artifactual noise. Thus the filtering of slow signal by derivatives combined with the scale-invariant properties of cones make the classifier robust to various sources of artifact and within-subject variability. Lastly, we will show that in addition to robustness to noise, our conic method is especially tuned towards task-related signals as opposed to contaminant spontaneous activity.

### 3.2. Sensitivity to fast-slow interactions

In general, the current literature suggests that very slow changes in the EEG amplitude are particularly prone to drift and motion artifact (i.e. [31]). In contrast, increasing evidence suggests that fast neural activity is associated with task performance ([32]). However, this does not necessarily indicate that slow components are always artifactual, but simply that fast components, when present, may be more informative. There are also many cases in which neural activity manifests meaningful fast and slow interactions ([33], [34], [35]). Thus, a method is needed that 1) can analyze slow signals when fast signals are weak/absent, 2) penalizes slow signals in the presence of faster ones, and 3) prioritizes slow components that exhibit slow-fast coupling over those that don’t. The derivative covariance matrix exhibits all of these properties, especially when the components have similar variance (related to spectral power). Formally, consider two multivariate processes: *X*(*t*), *Y*(*t*): ℝ^+^ → ℝ^*n*^ and class labels *i*(*t*) ∈ {*k*}. As before, we will denote the class-dependent derivative covariances as 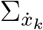 and 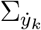. Now consider a second, slower process *B*(*t*) and a constant 0 < *c* < 1 defined such that for any class-invariant interval *τ* (as defined above) and non-negative scalars *t*, *ϵ*: [*t*, *t* + *ϵ*] ⊆ *τ* we have *B*(*t* + *cϵ*) − *B*(*t*) = *Y*(*t* + *ϵ*) − *Y*(*t*). In other words, *B*(*t*) evolves *c* times the speed of *Y*(*t*). The covariance matrix for the combined signal *S*(*t*):= *X*(*t*) + *B*(*t*) is:

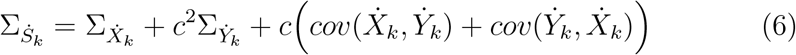

The notation *cov*(*A,B*) indicates the covariance between two multivariate processes, whereas we have previously indicated single-process covariance Σ_*A*_ as shorthand for *cov*(*A, A*). Clearly *c*^2^ < *c* and if 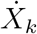,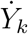 are independent, 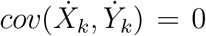. When 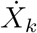 is sufficiently small (with the frequency profile held constant), the classifier’s inner term 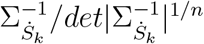 approaches 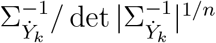 which satisfies the first condition: sensitivity to slow components in the absence of fast ones. Likewise as *c* becomes small (for 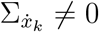), the classifier’s inner term is biased towards the higher frequency *X* term even if the components have similar amplitude thus satisfying our second condition. Lastly, when *X*, *Y* have amplitude, the classifier is more sensitive to the interaction between fast and slow components via 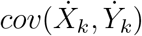 than those that are independent of *X* (hence restricted to 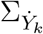) as *c*^2^ < *c*. Thus our classifier satisfies all three conditions to balance the influence of fast/slow signal components in a manner that emphasizes task-related components. If all three scenarios are mixed throughout a single class, direct calculation of the covariance based upon 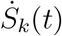 will be biased in favor of periods featuring fast components due to their greater derivative magnitude. Again, this bias can be removed by normalizing individual data-points of the derivative time series (using any norm). This step will change the calculated covariance for each class, but will not affect the data’s classifiability as 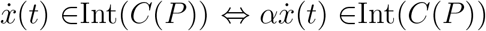 for any nonnegative *α*. The classifier’s performance regarding the three criteria is exemplified by the continued sensitivity in the bursting network example (Figure 2) which featured fast-slow coupling of downstream/upstream dynamics and within-cell fast-slow coupling during burst activity ([24]).

### 3.3. Continuous-Time Classification Reveals the Evolution of Cognitive States

As illustrated in Figure 4 the conic approach not only provides an accurate prediction for the trial, but also generates a prediction for each time point. Together, these predictions form a time series describing the evolution of the system between states with high temporal resolution. The key step of our conic classification technique is capturing dynamic properties by converting to the derivative time series. Previous dynamic approaches have generated state time-series using parameter estimates at fixed or sliding windows ([15],[16],[36],[37]). In contrast, the current approach generates a time series of individual predictions from a single classifier. Users have additional flexibility in choosing the desired resolution via smoothing etc., or application of secondary classification techniques to the cone-generated time series for higher-order analyses.

### 3.4. Extension to hemodynamic signals

Presently, we have only presented results of conic decoding for data with relatively high sampling rates, namely neuronal simulations and EEG. However, the potential exists for applications involving much slower modalities provided the events of interest are observable on a similarly long timescale. Task-related dynamics have long been observed in the BOLD signal (e.g. [38]) and more recent studies have also found dynamic relations between regions (e.g. [39],[40],[41]). Although the low sampling rate of fMRI is nonideal for derivative estimation, the aforementioned work suggests BOLD dynamics may be observable over longer intervals. The temporal derivative of interpolated data might then prove a suitable, smoother proxy. As such, the proposed method of conic classification has potential application to fMRI studies of temporally extended cognitive states.

### 3.5. Summary

We applied conic decoding to two sets of simulated data and one dataset of task-EEG to illustrate its usefulness in both supervised and unsupervised classification of system states. In the first simulation we simulated a system which under which two states were intractable in the original and spectral spaces, but immediately separated in derivative space. While the manner in which the system traversed state-space differed between cases, the overall range and distribution over the space remained constant, as might be expected in EEG/MEG recordings.

In our second example we modeled the case of a scientist attempting to determine the evolution of up-stream network states based upon the recorded voltages of three downstream cells. The cells used realistic phenomenological models with intrinsic bursting [24]) and synaptic dynamics. Despite using a simulated system, this case is of high empirical relevance. By varying system parameters we demonstrated that the advantage of our conic approach is greatest over spectral methods when couplings are relatively large or window lengths are relatively small, as might occur for a system rapidly shifting between states. In this specific example, the conic techniques did exhibit greater effect of measurement noise than spectral classification (of slow components), but neither system was sensitive to the intrinsic system noise. Thus the conic method suffers with extreme values of idealized, instantaneous white noise which impairs derivative estimation. However, the conic method is also invariant to many specific sorts of artifactual or recording noise, as discussed above. As such, the conic method has some inherent robustness to noise, despite the well-documented properties of derivatives in amplifying high-frequency noise. In scenarios where high-frequency is a factor, more sophisticated derivate estimation techniques, such as the total variational derivative ([42]) may be of use.

For the covert orienting (EEG) dataset, the method performed as-good or better than previously-published frequency-based decoding ([26]) at the group level (Figure 3A,B), but the negatively-trending correlation in performance at the individual level (Figure 3C)suggests that these approaches are sensitive to different signal properties and might benefit from being used together. By remapping the conic weights to spatial locations, we showed how our method reveals retinotopic organization of posterior channels during covert attention. These findings demonstrate that the conic decoding method allows simultaneous tracking of multiple cognitive states that may be particularly advantageous for the non-oscillatory signal components that hinder spectral decoding.

## 4. Materials and Methods

### 4.1. Derivation of the Conic Classification Criteria (2)

To derive our equation for conic classification we assume that the distributions for the two classes are elliptical in the sense that they may be described in terms of 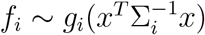. Unlike the traditional definition of an elliptic function such as those described in standard quadratic discriminant analysis we do not assume that the function *f* is non-increasing. In fact, this must not be the case from the perspective of autonomous dynamical systems as the origin/‘mean’ in derivative space corresponds to a fixed point and should thus have low density in the case of neural signals which are highly transient. Thus we assume nothing regarding the distributions *f*_*i*_ save that they are non-degenerate. Instead of restricting the distribution, we restrict our set of classifiers to be scale-invariant (conic) and thus we may instead consider the distribution projections on the surface of the (*n* − 1)-dimensional sphere (𝒮^*n*−1^) without loss of generality. What follows involves deriving the probability density of an elliptic distribution onto ∂𝒮^*n*−1^. These densitites immediately lead to the derivation of a maximum likelihood classifier of the form (2).

We impose zero density at the origin as it has no direction and hence is never classifiable. From a practical standpoint this assumption should be insignificant for continuous valued data with a finite number of data-points. Our treatment of the general elliptic case is in essence the same as the Gaussian case with mean zero long-considered in directional statistics [43]. We then apply simple monotone transformations to the distribution to ease computation in high dimensional settings.

**Proposition:** Consider a set of elliptical probability distributions {*f*}_*i*=1…*m*_ over ℝ^*n*^ with all means equal zero and *f*_*i*_(0) = 0 ∀*i* ∈ {1‥*m*}. Denote the corresponding covariance matrices for each elliptical distribution as {Σ}_*i*=1…*m*_, and a random variable *X* valued over ℝ^*n*^. Define the projection operator Π_*s*_: ℝ^*n*^ \ {0} → ∂S^*n*−1^ and its derived distributions (*h*_*i*_) as follows:

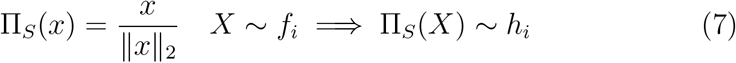

Then the following two assignment functions are equivalent:

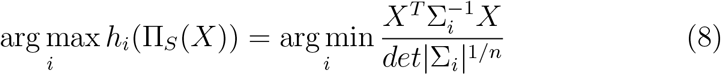

*Proof.* Consider the elliptic distribution *f*_*i*_ and covariance matrix Σ_*i*_. Thus, by the definition of an elliptic function, there is a function *g*_*i*_: ℝ → ℝ and a functional *κ*: *L*^1^ → ℝ such that:

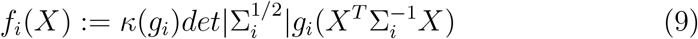

Here *κ*_*gi*_ denotes the scaling function which depends upon *g*_*i*_ but not Σ_*i*_.

The distribution for Π_*s*_(*x*) is the projection of *f*_*i*_ onto ∂𝒮^*n*−1^. In order to perform this radial projection we convert to spherical coordinates *r*(*x*) ∈ ℝ^+^ and *ϕ*(*x*) ∈ ℝ^*n*−1^ with radial variable *r*(*x*):= ‖*x*‖_2_ and angular vector *ϕ*(*x*) (for any full rank angular basis). We define the function Θ_*i*_ to represent the portion of the quadratic term, which depends upon the orientation of *x* (i.e., *ϕ*(*x*)), but not its magnitude:

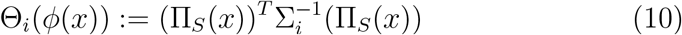

The distribution of Π_*S*_(*x*) ∈ ∂𝒮^*n*−1^ is found by integrating Θ_*i*_ along the radial component of *x*; *r*:= ‖*x*‖_2_:

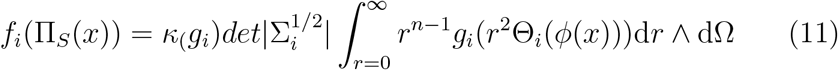

In which dΩ is the volume element of the (*n* − 1)-spherical shell. Using a simple change of variables (*u* = *r*^2^Θ(*ϕ*)) we remove the angular component Θ(*ϕ*):

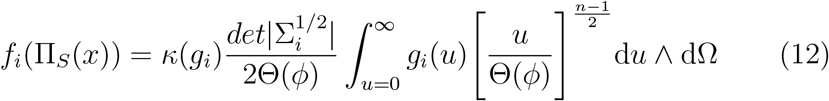

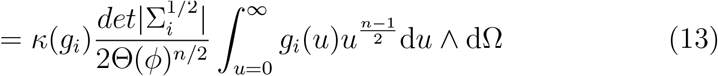

We will refer to the remaining integral as *ξ*(*g*_*i*_). Now consider a second distribution *f*_0_ characterized by a separate scalar function *g*_0_ ≠ *g*_*i*_, but identical covariance: Σ_0_ = Σ_*i*_;, hence Θ_0_(*ϕ*(*x*))=Θ_1_(*ϕ*(*x*)). The distributions may then be written:

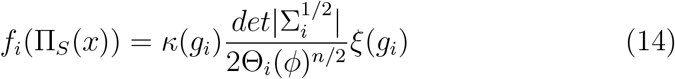

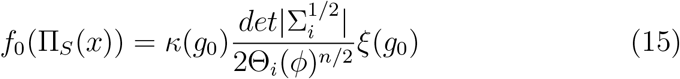

As both functions are probability distributions they must sum to 1. As the distributions are defined over ∂𝒮^*n*−1^ they are integrated over dΩ. Factoring out the scalar terms produces integrals dependent only upon the angular component Θ_*i*_(*ϕ*).

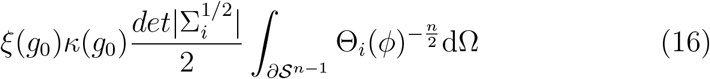

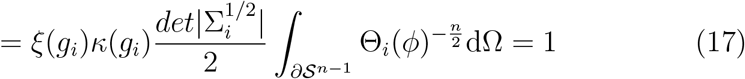

Thus there must be a constant *η*_*n*_ related to the geometry of the *n* − 1-sphere, such that *ξ*(*g*_*j*_)*κ*(*g*_*j*_) = *η*_*n*_ for any appropriate function *g*_*j*_. Dividing *f*_*i*_ by *η*_*n*_ and raising the result to the −2/*n* completes the proof:

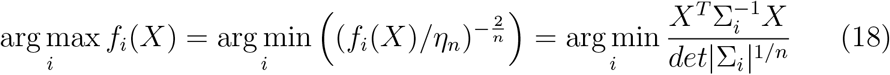

which amounts to our classifier (2).

To recover a full maximum likelihood classification which considers initial population proportions (*p*_*i*_) we simply multiply the derived probability density proportion of the probability density *f*_*i*_(*X*) by *p*_*i*_ before raising solutions to −2/*n* hence:

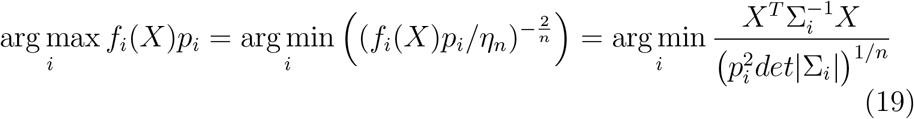

### 4.2. Computational Considerations and Limitations

A key computational step is calculation of the determinant. To avoid numerical instability associated with matrix inversion, we calculate the determinants via:

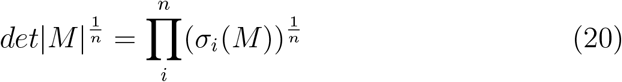

where σ_{1…n}_ indicate the singular values of *M*. Nevertheless, retrieving the eigenvalues may be computationally expensive (*O*(*n*^*k*>2^)) as compared with linear classification algorithms and this remains a limitation of the proposed method. However, we have used the default MATLAB eigenvalue algorithm for conic classification of 2,000 dimensional data on a midrange laptop (x64, dual Intel(R) Core(TM) i7-6500U @ 2.50GHz).

Another limitation of the current method concerns when the covariance matrices are ill-conditioned. In some potential applications (e.g. fMRI) the dimension of the data (number of channels) may exceed the number of datapoints per class. The simplest approach to this issue is dimensionality reduction or to (artificially) increase the number of datapoints through interpolation. However, an alternative method is to retain the original number of dimensions and use sparse-estimation for the population inverse matrix. Previous authors (e.g. [44]) have developed computationally efficient methods to perform this calculation. Many of these approaches rely upon LASSO or regularization methods so their accuracy relies upon at least some channels being independent.

### 4.3. Simulation 1

In Figure 1 we used a sigmoidal 3-neuron recurrent network to illustrate conic dynamics in a low dimensional space. The connection scheme is depicted schematically in Fig 1A,B and features two coupled negative feedback loops. A common hub (neuron 3) inhibits excitatory cells 1,2. Connections between neurons 2,3 were stronger than those with neuron 1. The model was as follows:

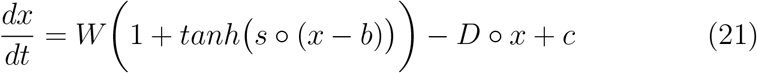

We use the notation o to indicate the Hadamard product (element-wise multiplication). We simulated this network twice using the same slope (*s*) and connection weights (*W*) for each case:

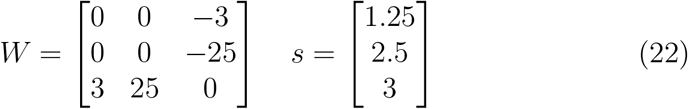

The two cases differed, however, in the values for decay (*D*), baseline (*c*), and threshold (*b*):

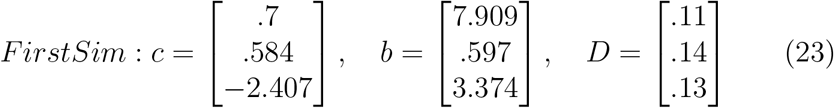

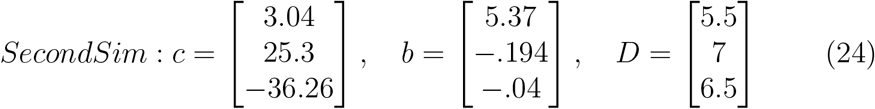

Simulations were performed with Euler integration *dt* = .0005 and a run time of *t* = 3 for each iteration and then down-sampled by a factor of 15. For each case 150 iterations were performed with starting positions selected pseudo-randomly. Segments of each time-series were also pseudorandomly selected using a random box-function (*B*(*t*). The box function was formed by choosing 200 box-lengths distributed according to (1 + *r*)^2^ with r distributed according to the standard normal distribution (*r N*(0,1)). The resulting box-lengths were then rescaled so that they summed to the total window size (*t*=3). The amplitude for each box was generated from a standard normal distribution and the inclusion rule for displaying a point x(t) was *B*(*t*) > 1.5. Thus, roughly 6.9% of the total points were displayed for each trajectory/initial condition.

All simulations were carried out in MATLAB2016b.

### 4.4. Simulation 2

For this simulation we considered a network of 4 bursting Hindmarsh-Rose neurons with single-exponential synapses. A chaotic Lorenz attractor provides input to the network as a representation of upstream network activity. The Lorenz attractor was given a standard parameterization with the addition of a Gaussian white-noise term (*η*_*i*_) with standard deviation *ϵ* (“Internal Noise”):

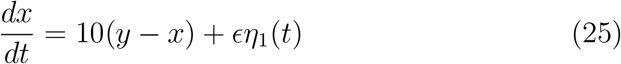

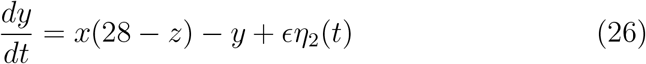

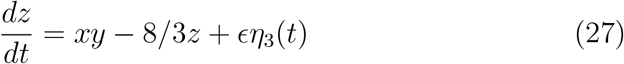

The Lorenz attractor was Euler integrated with dt=.0025 for duration t=50. Each time point was then upsampled (repeated) by a factor of 100 to ensure that the system did not fluctuate too quickly. The upstream system was considered ‘on’ when the first Lorenz variable (x) was greater than 0. Initial conditions for each iteration were normally distributed with mean [−10 −10 27] and standard deviation .25.

The output current from the Lorenz attractor was a binary function of the first Lorenz variable’s sign:

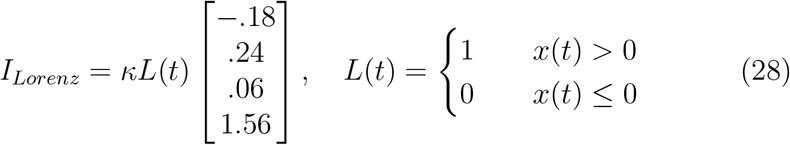

The term *κ* is a scaling factor (“Stimulus Strength”) which we manipulated. Each neuron’s internal dynamics were governed by the standard Hindmarsh-Rose equations [24]:

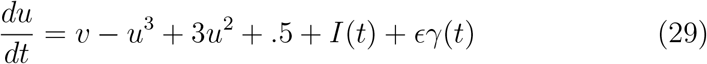

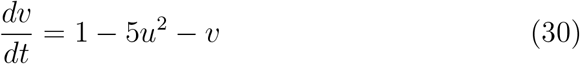

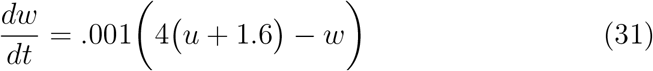

Here *I*(*t*) denotes the combined input to each cell and γ(*t*) denotes a Gaussian white-noise process with standard deviation *ϵ* (“Internal Noise”). Input consisted of a constant baseline level of input *I*_*base*_, the synaptic currents *I*_*Syn*_ and the currents from the Lorenz attractor *I*_*Lorenz*_. The synaptic currents were the product of a synaptic weight and a single dynamic synapse variable per neuron.

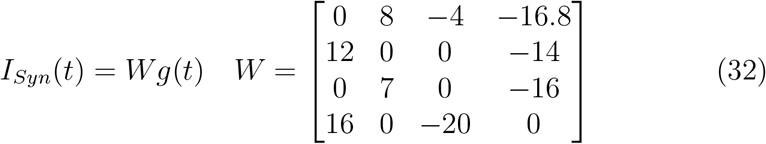

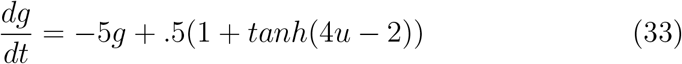

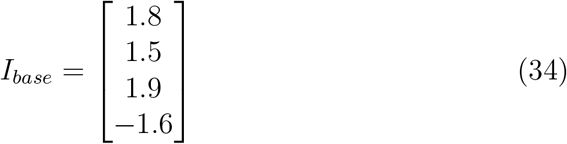

The Hindmarsh-Rose system was Euler integrated with step size .05 and end time t=5,000. Initial conditions for all neuron variables (u,v,w) were drawn from a standard normal Gaussian distribution. All synapses were given initial condition zero. The initialization period from t=0 to t=1,000 was removed to allow the system time to stabilize. The variables of interest consisted of the voltages (u) for the first three neurons only, which were down-sampled by a factor of 10. Gaussian noise was added with standard deviation *ξ* (“Measurement Noise”). We performed blind classification with clustering to assign system states (Lorenz input on/off) based upon the first three neurons’ voltages. We performed classification using the covariance of the normalized derivative time series (Δ*x*_*t*_/ ‖ Δ*x*_*t*_ ‖_2_) for each time bin as input to the k-means algorithm. Clustering for covariances was performed using k-means for 2 clusters for the upper triangle of the covariance (six elements). Similar clustering for spectral analyses were based upon 2-cluster k-means for the vector composed of the separate real and imaginary components of the spectrum evaluated at 6 equi-spaced points spanning half the smallest window size. The conic decoding was then performed using the two centroids derived from the k-means clustering as the covariance matrix for each class. The prediction weights for all points within a window were averaged to assign a class to the full window. For spectral clustering, labels were assigned directly based upon the centroid cluster to which each bin belonged. As the analysis involved blind clustering, cluster labels were arbitrary. In assigning cluster numbers to system states (i.e. does cluster 1 or 2 indicate input ‘on’) we used the assignment that provided maximal accuracy for each simulated trial. As such, the minimum accuracy for each trial is .5 and the expected accuracy is greater than .5 using this assignment. Therefore, we derive the null distribution for this case to evaluate bias. As the accuracy for a random assignment (random mapping between cluster number and ‘on’/‘off’ state) is uniform *U*[0,1], their mean is Bates distributed. For larger numbers of samples (as was the case for each trial), the Bates distribution for mean .5 over [0,1] (as there are two classes) limits to the Gaussian *N*(.5,1/(12*n*)). Thus the chance accuracy for random label assignment limits to *N*(.5,1/(12*n*)). For our biased case (using the assignment that provides greatest accuracy), the assignment function is *max*(*μ*(*A*), 1 − *μ*(*A*)) with A the accuracy obtained by arbitrary assignment. For several hundred samples, as was our case for each trial, the expected values of this distribution is essentially .5 (limiting to 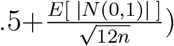.

To analyze the effect of parameterization we manipulated each parameter (intrinsic/measurement noise, stimulus strength, bin length) independently while keeping all others at their median value. There were a total of 136 trials per condition (parameter level) and 9 equi-spaced conditions per variable. As demonstrated in [2D], long run times ensured that despite the obvious noise/chaos within each trial [2B] standard deviations for decoding accuracy were relatively small. Combined with the large sampe size, these factors generated highly significant results for nearly every statistical comparison. As such we focused upon displaying performance. Note that the shading in [2D] indicates standard deviation as opposed to standard error (which would not be visible).

#### Experiment 3

The data and task were obtained from an open repository associated with ([26],[14]). Briefly, during the task, a multicolored hexagonal cue was presented for 200ms and participants were instructed to focus upon the edge corresponding to a given color. The orientation of this edge predicted in which of 6 radially located discs the upcoming target would be displayed after a delay of 500 – 2000ms. In 80% of trials the target appeared in the cued location. Targets were presented for 200ms after which participants had to indicate the target’s shape (either ‘+’ or ‘x’). The dataset from these studies is publicly available (BNCI-Horizon-2020.eu) and consists of eight subjects’ pre-processed recordings for 60 EEG channels, 2 EOG channels. Data was recorded at 1KHz but the published dataset has been down-sampled to 200 Hz. Our only further modification to the data was a single wavelet reconstruction to estimate derivatives given noisy data. For this step, we simply used a 1-D Daubechies 7-tap decomposition and level 3 reconstruction, although more specialized methods exist ([45]). The resultant signal was then converted to its difference (derivative) time series. Each data-point was scaled to have unit 2-norm; however, this step does not influence classifiability due to the scale invariance of cones. Rather this procedure only affects how the cone is parameterized by reducing the influence of outliers in estimating covariances. Analyses were repeated without normalization which produced near-identical results (within ±1% change in mean accuracy). Statistical analyses between previous and current results are based upon paired t-tests for each case. The significance of pair-wise accuracy was assessed with 1-sample t-tests for accuracy greater than chance (50%) and Bonferoni corrections which mimicked the original analyses with this dataset ([14]). For the 15 possible pairwise-comparisons this approach resulted in corrected significance thresholds of roughly 65% for *p* < .05.

Plots in Figure 4 display conic classification of subject 3’s data for the lower right vs. upper left contrast for the 21^st^, 24^th^, 27^th^, and 30^th^ trial of each of the two locations. Time series in 4B are the concatenated time series of the largest eigencomponent for each location across the four classification periods each. In Figure 4C the small blue circles denote the start of a new trial’s classification period and the black squares denote the end of the period.

#### 4.4.1. Generating Spatial Maps

All spatial maps displayed were generated using the group-averaged covariances of the derivative for each class. Data in 5 is displayed from a posterior coronal view (as indicated in center). For each location we considered the cone generated by the contrast location “x” vs. the hemifield opposite “x” (the average covariance across 3 locations). The method in which we visualized the conic matrix is by conversion to a vector which forms a static head-map. As with functional connectivity, the more natural way to visualize the matrix is as a weighted graph between nodes, however, we chose to use a headmap to better visualize the evident retinotopic organization. To do this, we defined each channels contribution to the conic matrix as the sum of its squared contributions to each eigenvector *v*_*j*_ weighted by the corresponding eigenvalue λ*j*. Thus the *i*^*th*^ channel’s weight (*w*_*i*_ was:

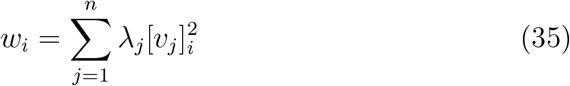

Smooth maps were created by interpolating the channel-wise measurements after coronal projection. Linear triangulation-based interpolation was performed with the built-in MATLAB function ‘griddata’.

#### 4.4.2. Eigenlengths and Eigencomponents

In the two class case (labels= ±1), the conic boundary corresponds to 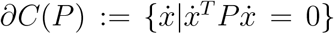 with P the weighted inverse of covariance matrices:

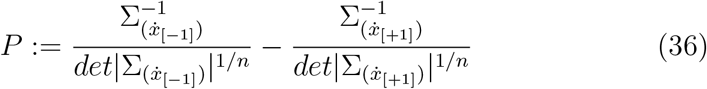

The class assignment function is 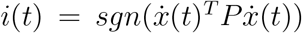. As P is real symmetric, it has a spectral decomposition *UDU*^*H*^. Splitting the diagonal matrix of eigenvalues (*D*) into positive and negative components: *D* = *D*^+^ − *D*^−^ produces:

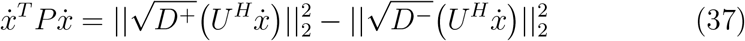

As the right side of Equation 37 involves the difference of two norms, we refer to them as the positive (with *D*^+^) and negative (with *D*^−^) eigenlengths. These eigenlengths are useful for generating a 2-dimensional projection of the data by which to compare separability as in Figure 4C. Similarly we generate “eigencomponents” of the system by simply rotating it into the cone’s coordinate-axes 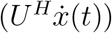. We call the eigencomponents positive or negative based upon the sign of the corresponding eigenvalue in *D* and these may be used to linearly decompose the signal into class-sensitive components as in Figure 4A,B. In general, the magnitude of each component’s eigenvalue is related to how tuned that component is for a specific class.

## Acknowledgments

MS was funded by NSF-DGE-1143954 from the US National Science Foundation. TB acknowledges R37 MH066078 from the US National Institute of Health. SC holds a Career Award at the Scientific Interface from the Burroughs-Wellcome Fund. Portions of this work were supported by AFOSR 15RT0189, NSF ECCS 1509342 and NSF CMMI 1537015, from the US Air Force Office of Scientific Research and the US National Science Foundation, respectively.

